# KCC2-dependent Steady-state Intracellular Chloride Concentration and pH in Cortical Layer 2/3 Neurons of Anesthetized and Awake Mice

**DOI:** 10.1101/234476

**Authors:** Juan Carlos Boffi, Johannes Knabbe, Michaela Kaiser, Thomas Kuner

## Abstract

Neuronal intracellular Cl^-^ concentration ([Cl^-^]_i_) influences a wide range of processes such as neuronal inhibition, membrane potential dynamics, intracellular pH (pH_i_) or cell volume. Up to date, neuronal [Cl^-^]_i_ has predominantly been studied in model systems of reduced complexity. Here, we implemented the genetically encoded ratiometric Cl^-^ indicator Superclomeleon (SCLM) to estimate the steady-state [Cl^-^]_i_ in cortical neurons from anesthetized and awake mice using 2-photon microscopy. Additionally, we implemented superecliptic pHluorin as a ratiometric sensor to estimate the intracellular steady-state pH (pH_i_) of mouse cortical neurons *in vivo*. We estimated an average resting [Cl^-^]_i_ of 6 ± 2 mM with no evidence of subcellular gradients in the proximal somato-dendritic domain and an average somatic pHi of 7.1 ± 0.1. Neither [Cl^-^]_i_ nor pH_i_ were affected by isoflurane anesthesia. We deleted the cation-Cl^-^ co-transporter KCC2 in single identified neurons of adult mice and found an increase of [Cl^-^]_i_ to approximately 26 ± 8 mM, demonstrating that under *in vivo* conditions KCC2 produces low [Cl^-^]_i_ in adult mouse neurons. In summary, neurons of the brain of awake adult mice exhibit a low and evenly distributed [Cl^-^]_i_ in the proximal somato-dendritic compartment that is independent of anesthesia and requires KCC2 expression for its maintenance.

## Introduction

Neuronal [Cl^-^]_i_ is of crucial importance for processes such as inhibitory neurotransmission, maintenance of the resting membrane potential, regulation of intracellular pH and cell volume (Kaila et al., 2014; Doyon et al., 2016). This work is an attempt to gain insight into mammalian neuronal [Cl^-^]_i_ *in vivo* and thus contribute to the current knowledge in this field, which stems mainly from *in vitro* experiments. Although invaluable ground breaking work about neuronal [Cl^-^]_i_ has been produced *in vitro* (Delpire, 2000; Ben-Ari, 2002; Kaila et al., 2014), many factors inherent to routine *in vitro* preparations such as brain slices may influence [Cl^-^]_i_ regulation (Dzhala et al., 2012). Recent pioneering work achieved *in vivo* measurements of bulk tissue Cl^-^ dynamics (Wimmer et al., 2015; Berglund et al., 2016; Wells et al., 2016), but presently we know very little about the *in vivo* steady-state neuronal [Cl^-^]i and the factors affecting it (Bregestovski et al., 2009; Arosio and Ratto, 2014; Kaila et al., 2014; Doyon et al., 2016). Clomeleon was the first genetically encoded sensor to measure intracellular Cl^-^ and consists of a CFP-YFP pair in which the YFP FRET-acceptor is the Cl^-^ sensitive variant Topaz (Kuner and Augustine, 2000). However, the original version of Clomeleon has a Cl^-^ affinity of ∼160 mM that is out of the expected physiological neuronal [Cl^-^]_i_ and a limited signal to noise ratio (Grimley et al., 2013). This led to the development of a second generation of promising genetically encoded Cl^-^ indicators with improved affinity and signal to noise ratio that could be used for *in vivo* applications, such as ClopHensor (Arosio et al., 2010) and Superclomeleon (Grimley et al., 2013) (SCLM).

The affinity of most Cl^-^ sensitive fluorescent proteins for Cl^-^ is highly dependent on pH (Arosio et al., 2010; Grimley et al., 2013; Arosio and Ratto, 2014), which demands monitoring the cellular pH to validate the Cl^-^ readings made with these indicators. The design of ClopHensor-based sensors allows for simultaneous ratiometric pH and Cl^-^ measurements, but at the expense of being an excitation and emission ratiometric sensor. This hampers the *in vivo* implementation of ClopHensor-based sensors as its different excitation and emission wavelengths might be differentially scattered throughout the tissue at increasing imaging depths. While the present work was under review, a study by Sulis Sato et al. (2017) reported simultaneous *in vivo* optical estimations of neuronal Cl^-^ and pH in anesthetized mice using an optimized version of ClopHensor. The strong depth-dependent differential light scattering of the different wavelengths was compensated for by an offline correction algorithm. SCLM is not exempt of differential depth-dependent light scattering, but as it is an emission only ratiometric sensor instead, it might produce measurements less influenced by light scattering. This is at the expense of the need to monitor intracellular pH by other means.

Although neuronal pH has been studied *in vivo* using techniques such as nuclear magnetic resonance (Vorstrup et al., 1989), up to date mammalian neuronal pHi has not been studied with cellular resolution *in vivo*. The use of a genetically encoded ratiometric fluorescent indicator is again a promising alternative to achieve this. Recently, the implementation of superecliptic (SE) pHluorin (Miesenbock et al., 1998) as an emission ratiometric pH indicator to monitor neuronal pH_i_ has been reported in flies (Rossano et al., 2013). This suggests a strategy combining ratiometric SCLM and SE-pHluorin recordings as a promising alternative to study neuronal steady-state Cl^-^ and pH *in vivo* using 2-photon microscopy.

Thus, to contribute to the on-going *in vivo* implementation of Cl^-^ imaging we established *in vivo* 2-photon microscopy in mice expressing SCLM or SE-pHluorin and explored the strengths and caveats of this ratiometric *in vivo* imaging approach. To gain insight into the factors affecting neuronal intracellular steady-state Cl^-^ concentration, we compared [Cl^-^]_;_ and pH_;_ in anesthetized and awake mice as well as in single neurons lacking the cation-Cl^-^ co-transporter KCC2 (Delpire, 2000; Ben-Ari, 2002; Kaila et al., 2014). These results and approach will contribute to future studies of the role of Cl^-^ regulation in pathophysiological conditions such as epilepsy or schizophrenia *in vivo* (Kaila et al., 2014; Sullivan et al., 2015).

## Methods

### Ethics statement

This study was carried out in accordance with the European Communities Council Directive (86/609/EEC) to minimize animal pain or discomfort. All experiments were conducted following the German animal welfare guidelines specified in the TierSchG. The local animal care and use committee (Regierungspräsidium Karlsruhe of the state Baden-Württemberg) gave approval for the study under the number G183/15.

### Animals

Experiments were performed on wild type (WT) or homozygous knock-in mice bearing floxed KCC2 alleles. The mouse strains used were C57B16N/CR and KCC2^lox/lox^ (Seja et al., 2012; Godde et al., 2016). The latter mouse line was provided by Prof. Dr. Thomas Jensch (FMP/MDC, Berlin). All animals were 3-4 months of age by completion of data collection. Mice were housed up to three per cage and kept on a 12/12 h light/dark cycle. Food and water was available *ad libitum* except while the animals were restrained for imaging. Mice of either sex were used.

### Genetically encoded indicators

The SCLM variant (Grimley et al., 2013) of Clomeleon (Kuner and Augustine, 2000) was used for non-invasive imaging of [Cl^-^]_i_ in layer 2/3 (L2/3) pyramidal cells of motor cortex. SCLM consists of the Cl^-^-insensitive Cerulean variant of green fluorescent protein connected by a linker to the Cl^-^-sensitive Topaz variant. The Superecliptic variant of pHluorin (SE-pHluorin) (Miesenbock et al., 1998) was used for intracellular pH estimations. The Cerulean-Venus tandem pair and EGFP were used as variants less sensitive to Cl^-^ and pH of SCLM and SE-pHluorin respectively, to evaluate imaging depth dependent scattering effects on ratiometric imaging. In all cases, expression in L2/3 pyramidal cells of motor cortex was driven by the Synapsin minimal promoter using a Cre-dependent DIO cassette. Plasmids encoding SCLM, SE-pHluorin, Venus and Cerulean were kind gifts of Prof. Dr. George Augustine (DukeNUS Medical School), Prof. Dr. Gero Miesenböck (University of Oxford), Dr. Atsushi Miyawaki (RIKEN) and Prof. Dr. Dave Piston (Vanderbilt University) respectively.

### Calibration curves

All calibrations were done *in vitro,* as *in vivo* calibration is not feasible because of the lack of control of local conditions. Calibration curves were constructed as described elsewhere (Grimley et al., 2013). Briefly, [Cl^-^]_i_ and pH in cultured hippocampal neurons expressing SCLM or SE-pHluorin were matched to those of the extracellular saline by treatment with ionophores: 10 μM nigericin and 5 μM tributyltin acetate (Alfa Aesar). Cells were perfused in the presence of ionophores with calibration solutions of different [Cl^-^] or pH at a rate of 2 ml/min. High-[Cl^-^] calibration solution contained 105 mM KCl, 48 mM NaCl, 10 mM HEPES, 20 mM D(+)-glucose, 2 mM Na-EGTA, and 4 mM MgCh. Cl^-^ free calibration solution was composed of 10 mM HEPES, 20 mM D(+)-glucose, 48 mM Na-gluconate, 105 mM K-gluconate, 2 mM Na-EGTA, and 4 mM Mg(gluconate)_2_. Intermediate [Cl^-^] solutions were prepared by mixing these two solutions. The KF solution used to saturate SCLM contained 10 mM HEPES, 20 mM D(+)-glucose, 48 mM NaF, 105 mM KF, 2 mM Na-EGTA, and 4 mM Mg(gluconate)_2_. For SCLM calibration curves, all salines were adjusted to pH 7.10 or pH 7.45. For SE-pHluorin calibration curves a calibration solution of 5 mM Cl^-^ was titrated to different pH values. Image acquisition started after 10 min. of incubation with the first calibration solution containing the ionophores and subsequent incubations were 6 min. each. Only a maximum of 4 incubations were performed per coverslip to avoid deterioration of the culture. SCLM calibration curves were constructed by fitting scatter plots of FRET ratios vs. [Cl^-^] to the equation [Cl^-^] = K_d_((R_max_-R)/(R-R_min_))(C_f_/C_b_) where R is the YFP/CFP emission ratio, R_max_ and R_min_ are the maximum and minimum ratios, C_f_ and C_b_ are the absolute CFP fluorescence values in the Cl^-^ free and fully bound state respectively (Grimley et al., 2013). Average R_max_, R_min_, C_f_ and C_b_ were deteR_min_ed experimentally for the fitting equation. SE-pHluorin calibration curves were constructed by fitting scatter plots of 530 nm/480 nm emission ratios to the Boltzmann equation (Rossano et al., 2013).

### Viral vector expression

Adeno-associated viral particles (AAV2/1-2 serotype) driving Cre recombinase expression, Cre-dependent or -independent expression of SCLM, SE-pHluorin, Cerulean-Venus or EGFP, all under the control of the Synapsin minimal promoter, were prepared for stereotaxic injections (Schwenger and Kuner, 2010). To achieve sparse but bright labelling of neurons that permitted the identification of each cell’s projections, we co-injected AAVs encoding the Cre-dependent fluorescent proteins/sensors together with highly diluted (1/1000 to 1/5000) AAVs encoding nucleus-targeted Cre recombinase. Thus, due to the likelihood of the multiplicity of infection, very sparse but brightly labelled neurons could be imaged resolving their individual projections.

### Stereotaxic injection

For injection and craniectomy, mice were anesthetized by i.p. injection of a mixture of 40 μl fentanyl (1 mg/ml; Janssen), 160 μl midazolam (5 mg/ml; Hameln) and 60 μl medetomidin (1 mg/ml; Pfizer), dosed in 3.1 μl/g body weight, placed in a stereotaxic headholder (Kopf) and a circular craniectomy (∼7mm) centered at bregma was made with a dental drill. The dura was carefully removed over the area for later injection with fine forceps cautiously avoiding damaging blood vessels. AAV particles were injected at L2/3 M1 motor cortex using the following coordinates: in mm from the centre of the craniectomy, considered bregma, (x; y) = (1; 0). M1 cortex injections were performed using glass pipettes lowered to a depth of 300 μm to target L2/3. AAVs were injected using a syringe at a rate of ∼1 μl/hr. Volumes of virus of ∼0.2 μl were injected per spot. All mutant mice used for virus injection were homozygous for the floxed KCC2 allele. A round 6 mm diameter number 0 coverslip (cranial window) disinfected with 70% ethanol and a custom made round plastic holder crown surrounding it for head fixation were cemented to the skull using dental acrylic (Hager & Werken). We targeted L2/3 of M1 cortex due to the accessibility of this cortical layer for 2P imaging and the convenience of the location over M1 cortex for hosting a 6 mm round cranial window, which allowed for multiple AAV injection sites thus refining our technique to reduce the number of animals needed for our experiments in conformity with our ethics statement. The skin wound around the window was also closed with dental acrylic. Mice received an i.p. mixture of 120 μl naloxon (0.4 mg/ml; Inresa), 800 μl flumazenil (0.1 mg/ml; Fresenius Kabi) and 60 μl antipamezole (5 mg/ml; Pfizer) dosed in 6.2 μl/g body weight at the end of surgery to antagonize the anesthesia mix. Mice were given carprofen (Rimadyl, 5 mg/kg; Pfizer) as an analgesic immediately before surgery. Mice were single housed after surgery and typically imaged at least 28 days after virus injection and window implantation.

### Two-photon (2P) imaging

2P imaging was done on a TriM Scope II microscope (LaVision BioTec GmbH) equipped with a pulsed Ti:Sapphire laser (Chameleon; Coherent). The light source was tuned to 840 nm for SCLM and Cerulean-Venus measurements or 810 nm for SE-pHluorin and EGFP measurements. Imaging was performed with a 25X, 1.1 numerical aperture, apochromatic, 2 mm working distance, water immersion objective (Nikon, MRD77225) and emitted fluorescence was split with a 495 nm dichroic, then filtered with 540/40 nm and 480/40 nm emission filters for SCLM and Cerulean-Venus measurements or 530/70 nm and 480/40 nm for SE-pHluorin or EGFP measurements (Chroma). Fluorescence emission was collected with low-noise high-sensitivity photomultiplier tubes (PMTs; Hamamatsu, H7422-40-LV 5M).

### *In vivo* 2P imaging

For the imaging sessions of cranial window implanted mice, anesthesia was induced with 6% isoflurane (Baxter) in O_2_ and maintained at 0.8-1%. The anesthetized mouse was then head fixed on top of a rotating disc treadmill by a custom made head holder designed to fix a plastic holder crown surrounding the cranial window. This device was mounted onto the intravital stage of the 2P microscope. Fluorescence imaging in the anesthetized state was performed using line scanning. Typically, 16 bit frames of 491 μm^2^ and 1024 px^2^ resolution were acquired at a frame rate of 0.6 Hz and at 2 μm z steps. A full 3D stack spanning typically ∼300-350 μm in depth took ∼5-6 min. to acquire. Anesthetized imaging sessions lasted no longer than ∼30 min. After the imaging session in the anesthetized state, the animal was allowed to recover from anesthesia while still being head fixed at the microscope’s stage (usually it took the mice 2∼3 min. to show clear signs of being awake such as to start whisking, blinking and walking). Throughout the imaging session the animal’fluorescence intensity from each s breathing and behaviour was monitored by means of an infra-red camera mounted to the stage while keeping the stage in the dark. After 5 minutes from withdrawal of isoflurane, awake imaging was performed under the aforementioned settings. Awake sessions (since the withdrawal of anesthesia) lasted no longer than ∼15 min. After this the imaging session ended and the animal was returned to its cage.

### In vitro 2P imaging

Hippocampal cultures were prepared from E16 rats according to established techniques (Kuner and Augustine, 2000) Infection of neuronal cultures with AAVs delivering the different expression constructs was performed on (DIV) 7-8. After virus infection and 4-7 day incubation for fluorescent protein expression, coverslips with cultured cells were transferred to a custom made perfusion chamber with temperature control mounted on the intravital stage of a trimscope microscope (LaVision) equipped with a pulsed Ti:Sapphire laser (Chameleon; Coherent). All recordings were conducted at 35 ± 2 °C. Imaging was performed under the previously described settings. Except for calibration curves, hippocampal cultures were imaged in artificial cerebrospinal fluid (ACSF) gassed with carbogen (125 mM NaCl, 2.5 mM KCl, 25 mM NaHCO_3_, 1.25 NaH_2_PO_4_, 1 mM MgCl_2_, 2 mM CaCl_2_, 25 mM D(+)-glucose, pH = 7.4). All chemicals were purchased from Sigma-Aldrich, except otherwise noted.

### 2P imaging of ratiometric sensors

In a typical imaging session, the fluorescence emission collected at the shorter wavelength channel (cyan) was much weaker than the one from the longer wavelength channel (yellow/green). Hence, excitation laser power settings were optimized for cyan emission while avoiding saturation of the yellow/green emission channel, a compromise inherent to the dynamic range of ratiometric sensors based on CFP and YFP FRET pairs. To avoid “divided by background” ratio artefacts, we only analysed regions of interest (ROIs) that exceeded the emission intensity of background twofold. For the sake of consistency, we only analysed fluorescence ratios from cells with L2/3 pyramidal morphology, defined by the depth of the soma in the imaged volume, its size, shape and the presence of a marked apical dendrite (for simplicity referred to as pyramidal cells).

### Analysis of imaging data

Image analysis was performed using Fiji (Schindelin et al., 2012) and data analysis and statistics were performed using Prism 7 software (GraphPad). Regions of interest spanning the cell somas, dendrites, dendritic spines or a cell-free background area were determined manually. Mean fluorescence intensity from each ROI were subtracted with the mean fluorescence intensity from the corresponding background area and background corrected fluorescence intensity ratios were calculated. Due to the low intensity emission registered for Cerulean and SE-pHluorin at 480 nm, only ROIs in which this fluorescence intensity was twofold to that of the background were considered for further analysis.

### Statistical Analyses

Normality was tested using Kolmogorov-Smirnov, D’Agostino-Pearson and Shapiro-Wilk tests. For paired samples with normally distributed frequencies, statistical significance was evaluated using paired t-tests. Wilcoxon tests were used for pairwise comparisons in which one or both samples were not normally distributed. For unpaired comparisons between highly variable samples, the whole frequency distributions were compared using Kolmogorov-Smirnov tests. Statistical differences to a theoretical value was evaluated using Wilcoxon signed rank tests. Variances of samples with different sizes were compared with Levene's test. Correlation was evaluated using the Pearson correlation coefficient. A p < 0.05 was considered significant.

## Results

### *In vivo* imaging of SCLM-expressing layer 2/3 cortical pyramidal neurons in anesthetized mice

To explore the [Cl^-^]_i_ of neurons in the intact brain, we sparsely targeted SCLM expression to L2/3 of M1 cortex (Fig. 1A), typically obtaining one to ∼20 neurons expressing SCLM in the entire mouse brain (see methods for details). Using this criterion 31 pyramidal cells were analysed in dual color 3D stacks obtained from 5 head-fixed anesthetized WT mice. The Topaz component of SCLM gave rise to high fluorescence intensities even at depths of ∼300 μm from the brain surface (Fig.1A, B), yet Cerulean fluorescence was lower. At low [Cl^-^]_i_, Topaz effectively acts as an acceptor for fluorescence resonance energy transfer from Cerulean, hence limiting the overall brightness of Cerulean fluorescence emission. To illustrate the signal strength over background of each emission channel rather than the intensity differences between the channels, both channels were clipped for display (Fig. 1). In most neurons analysed, a section of the apical dendrite of L2/3 pyramidal cells, proximal to the soma, consistently produced bright fluorescence in both emission channels that allowed reliable analysis. Regions with signal to background ratios of less than twofold were excluded for calculating the 540 nm/480 nm emission ratio (SCLM ratio), with high ratios representing low [Cl^-^]_i_ (Fig. 1B).

**Figure 1:**
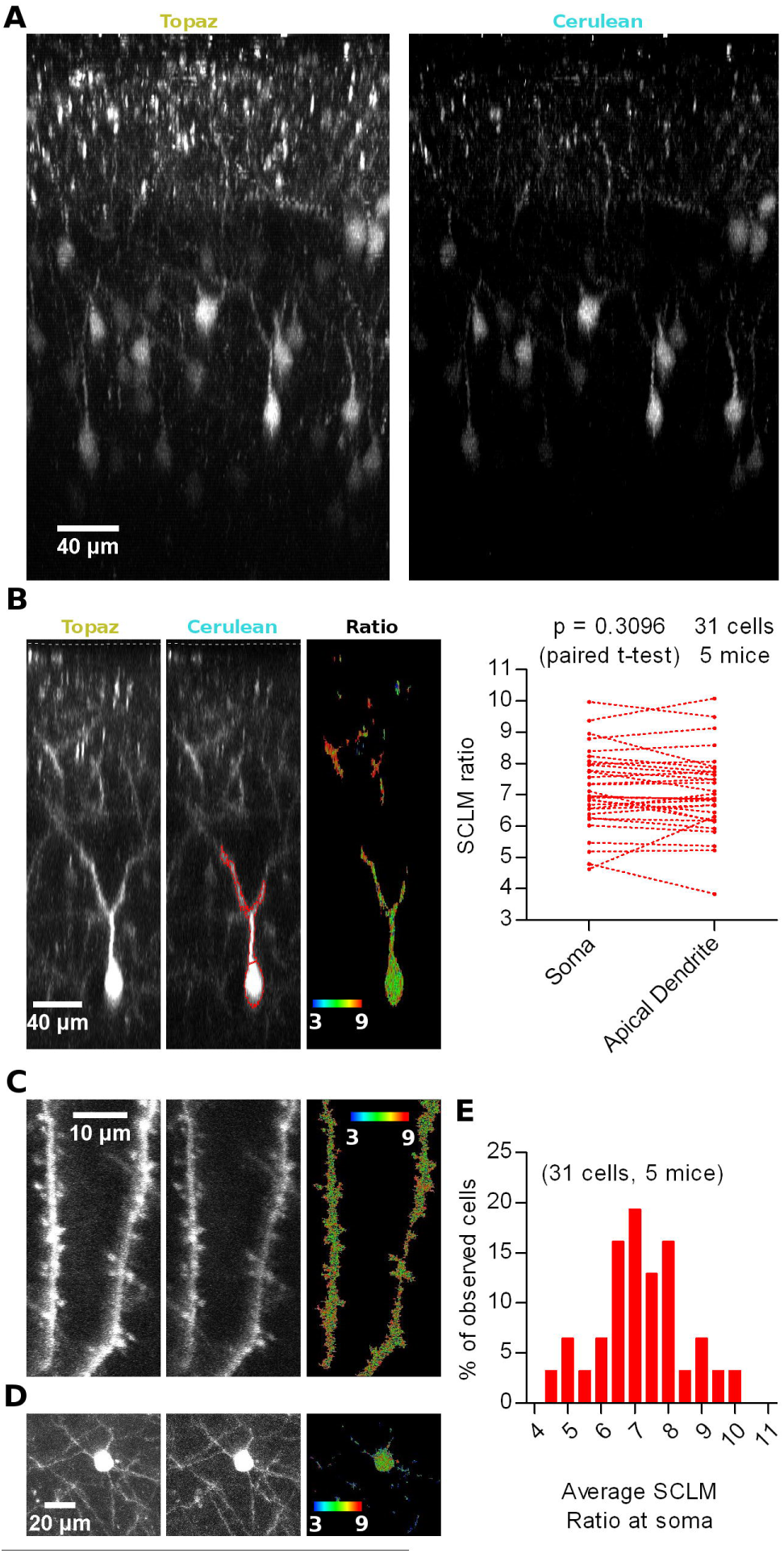
*In vivo* imaging of SCLM-expressing neurons in anesthetized mice. A) Representative maximal intensity projections of a vertically resliced data stack showing the SCLM Topaz and Cerulean signal of sparsely labelled L2/3 neurons of M1 cortex in an anesthetized mouse. Gray values were clipped to include > 90% of the total fluorescence signal: Topaz, 3557996; Cerulean, 247-1390 B) Left: Representative maximal intensity projections of a vertically resliced data stack showing the *in vivo* SCLM Topaz and Cerulean signals and their corresponding ratio of a L2/3 pyramidal neuron from M1 cortex in an anesthetized mouse. For display, gray values were clipped to include >90% of the total fluorescence signal: Topaz, 291-4536; Cerulean, 111-757. The displayed neuron's soma centroid is ∼270 μm deep from the surface of the brain (included in the acquired stacks and marked with a white segmented line on the Topaz and Cerulean panels). Representative somatic and dendritic ROIs enclosing areas with 2 fold signal to background emission intensities are displayed with red lines on the Cerulean panel. Right: Pairwise comparison of the average SCLM 540 nm/480 nm emission ratio from ROIs spanning the basal apical dendrite and soma of L2/3 pyramidal neurons from M1 cortex in the anesthetized state. C) Representative maximal intensity projections showing SCLM Topaz and Cerulean signals and their corresponding ratio of superficial spiny dendrites of L2/3 pyramidal neurons from M1 cortex in an anesthetized mouse. Gray values were clipped to include >90% of the total fluorescence signal: Topaz, 3063655; Cerulean, 103-774. D) Representative maximal intensity projections showing the *in vivo* SCLM Topaz and Cerulean signals and their corresponding ratio of the soma and basal dendrites of L2/3 pyramidal neurons from M1 cortex of an anesthetized mouse. Gray values were clipped to include >90% of the total fluorescence signal: Topaz, 90-2000; Cerulean, 96-295. E) Histogram showing the frequency distribution of the recorded average SCLM 540 nm/480 nm emission ratio from ROIs spanning the somas of L2/3 pyramidal neurons from M1 cortex in the anesthetized state.

A cell by cell pairwise comparison between the average SCLM ratio measured from a ROI spanning the soma or the apical dendrite of the 31 L2/3 pyramidal cells revealed no significant difference (p = 0.3096, paired t-test; Fig. 1B), suggesting that intracellular Cl^-^ levels are rather homogeneous throughout these two subcellular compartments. Distal dendritic and spine [Cl^-^]_i_ could be imaged in some neurons (Fig. 1C), nevertheless the lower signal to background ratio in these structures precluded a systematic analysis. Additionally, these few distal spiny dendritic stretches could not be traced back to a cell soma. Similarly, the Cerulean fluorescence at the basal dendrites of L2/3 pyramidal cells was too low to produce reliable ratios (Fig. 1D). These results suggest that [Cl^-^]_i_ would be evenly distributed within the proximal somato-dendritic domain up to ∼100 μm from the soma of L2/3 pyramidal neurons. Unfortunately, technical limitations do not allow measurements of [Cl^-^]_i_ in thin superficial dendrites and their spines as well as in thin basal dendrites deeper in the tissue.

We noticed that the fluorescence ratios differed substantially between individual neurons (Fig. 1E). The large range of reported SCLM ratios could be a consequence of several factors: (1) depth-dependent differential scattering of the two emission wavelengths, (2) local *in vivo* imaging conditions (i.e. number of large blood vessels within the imaging pathway or other conditions of optical inhomogeneities), (3) biological variability, i.e. neurons having different resting [Cl^-^]_i_ or pHi, and (4) other factors. These issues will be addressed in following sections.

In summary, *in vivo* SCLM imaging with cellular resolution is feasible within cellular compartments that produce sufficiently high fluorescence intensity in the Cerulean emission channel at illumination intensities that do not saturate the Topaz channel, i.e. somata and thick dendrites of neurons. Within the proximal somato-dendritic domain spanning ∼100 μm, [Cl^-^]_i_ would be evenly distributed with no detectable intracellular gradients in [Cl^-^]_i_.

### Depth-dependent differential scattering of cyan and yellow fluorescence?

An intrinsic issue of ratiometric sensors is that their different emission wavelengths could be differentially scattered by the tissue, hence photons of different wavelengths emitted by SCLM travelling through different amounts of brain tissue at different imaging depths could produce depth-dependent artificial ratios. To address this problem, we designed an environmentally insensitive SCLM variant by replacing the Cl^-^ and pH sensitive Topaz yellow fluorescent protein with the reduced environmental sensitivity YFP variant Venus (Nagai et al., 2002). The pK_a_ value of Venus (6, (Nagai et al., 2002)) is slightly lower than that of SCLM (6.4, (Grimley et al., 2013)) and the K_d_ for Cl^-^ of Venus (10 M, (Nagai et al., 2002)) is strongly increased. Thus, Venus can be considered insensitive to Cl^-^, yet slightly sensitive to pH. Cerulean is insensitive to pH in the range tested (pK_a_ = 4.7, (Shaner et al., 2005)) and its emission is not influenced by Cl^-^ (Kuner and Augustine, 2000; Grimley et al., 2013). Hence, we replaced Topaz in SCLM with Venus, obtaining Cerulean-Venus. Thus, depth-dependent ratio changes would indicate differential scattering, while an invariant ratio would argue for depth-independence.

We sparsely expressed Cerulean-Venus in L2/3 pyramidal neurons of the motor cortex, under the same settings used with SCLM. We then recorded emission ratios at somas and apical dendrites at different imaging depths and included the surface of the brain in the image stack to estimate the depth of the somas (Fig. 2A). *In vivo* 2P 3D stacks obtained from six head fixed anesthetized WT mice revealed that average somatic Cerulean-Venus ratios did not significantly correlate with the depth of the soma centroid (Pearson r = −0.01256, p = 0.9296, n = 52 cells; Fig. 2A, B). To further assess the influence of light scattering in our measurements, we studied the emission ratio along the longitudinal axis of the neurons from the base of the soma towards the surface of the cortex along the apical dendrite until the most superficial position possible (Fig. 2A, right panel). While individual cells had rather different ratios, we could not find systematic shifts of the ratio along the dendritic tree (Fig. 2C). We evaluated the correlation of the emission ratio *versus* the distance along the longitudinal axis for each cell and found both positive and negative correlations (Fig. 2D). The expected effect of scattering should manifest itself in only one sense of correlation, likely negative in this case due to higher scattering of shorter wavelengths. When analysing this correlation in a cell by cell basis we found that 67% of the cells (n = 52) displayed Pearson correlation coefficients between −0.3 and 0.3, which could be considered weak or no correlation. Only in 26% of the cells, we observed coefficients lower than −0.3 and in 7% higher than 0.3. Thus, we conclude that differential depth-dependend scattering, within the range covered by our experiments, does not contaminate the ratiometric readout.

**Figure 2:**
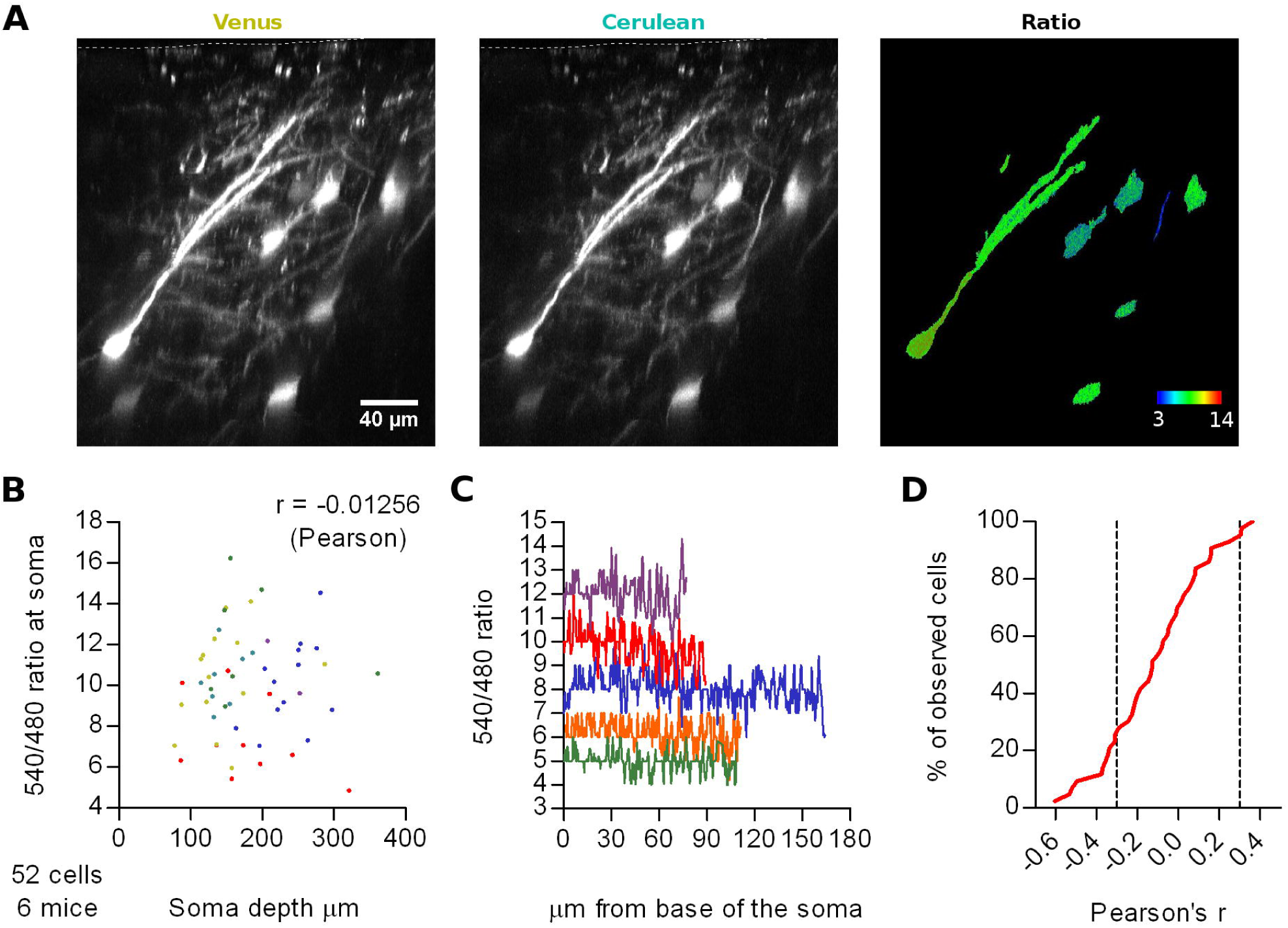
Effect of depth-dependent differential scattering. A) Representative maximal intensity projections of a vertically resliced data stack showing the Cerulean-Venus signal and ratios from somas at different depths and along an apical dendrite of L2/3 neurons from M1 cortex. The surface of the brain is marked with a white segmented line on the Venus and Cerulean panels. Gray values were adjusted as follows: Venus, 306-5000; Cerulean, 98-677. B) Average 540 nm/480 nm emission ratios from ROIs spanning the somas of these cells as a function of depth of the soma centroid. Each color represents data from one out of six mice. C) Representative traces of the 540 nm/480 nm ratio along the longitudinal axis of five neurons, from the base of the soma (0 μm) through the extent of the soma and apical dendrite until the last imaging plane in which the Cerulean signal had two-fold background intensity. Each color represents a different neuron. D) Cumulative frequency distribution of the Pearson's correlation coefficient evaluated for the 540 nm/480 nm emission ratio as a function of the distance from the base of the soma along the longitudinal axis of 43 neurons from 6 mice. Only cells with lengths bigger than 40 μm (∼twice the length of the soma) were considered, to span a representative stretch of imaging depths. Segmented lines mark 0.3 and −0.3 correlation coefficients.

Similar to the ratios recorded with SCLM, Cerulean-Venus ratios displayed a pronounced dispersion when comparing different cells (Fig. 2C). The observation that the ratios obtained with the environmentally much less sensitive Cerulean-Venus and SCLM are dispersed in a similar fashion, suggests that the observed dispersion in SCLM ratios is not due to variation in intracellular [Cl^-^]_i_ between cells. This point will be further elaborated below.

### *In vivo* ratiometric imaging of SE-pHluorin-expressing layer 2/3 cortical pyramidal neurons

SCLM presents a sensitivity to pH that cannot be ignored as pH significantly affects the K_d_ of Cl^-^ for SCLM (Grimley et al., 2013). To deteR_min_e the neuronal pH_i_, we imaged L2/3 neurons of the motor cortex expressing SE-pHluorin. SE-pHluorin has been successfully used as a ratiometric pH indicator *in vitro* using excitation and emission wavelengths similar to those used for SCLM (Rossano et al., 2013), facilitating the implementation of this sensor for ratiometric pH estimations through minimal modifications to our method. SE-pHluorin was excited at 810 nm and 530 nm and 480 nm emission filters were used to calculate the 530 nm/480 nm ratio as a measure of pH_i_ (see methods).

To deteRmine the average pH_i_ of the imaged cells, we constructed a calibration curve in SE-pHluorin-expressing hippocampal neuronal cultures by clamping pH_i_ to the extracellular saline by means of ionophore treatment at close to physiological temperature (35 ± 2°C). We fitted the data to the Boltzmann equation (r^2^ = 0.7009) obtaining a best fit V_50_ value of 7.49 ± 0.05 (Fig. 3A).

**Figure 3:**
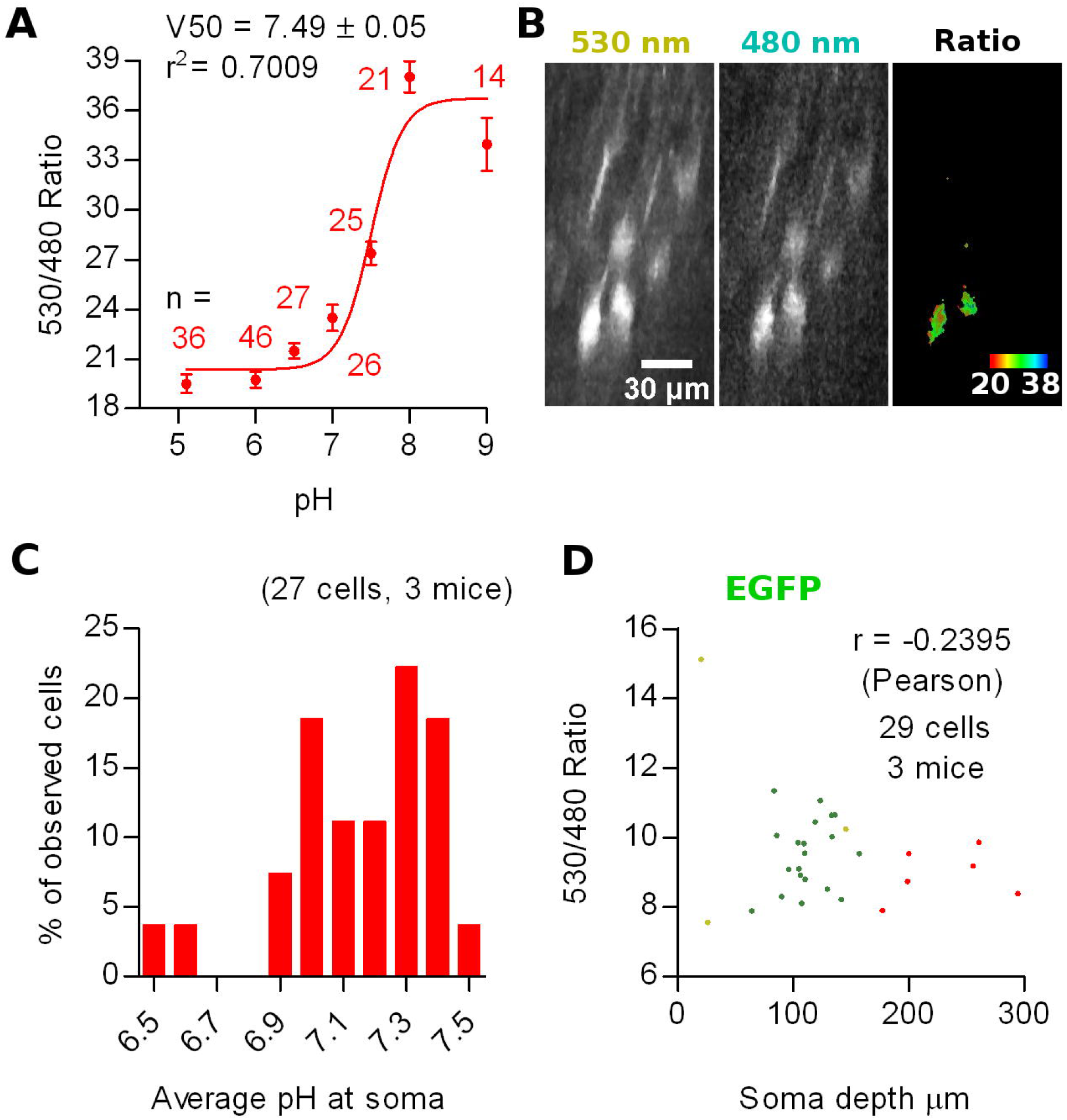
*In vivo* ratiometric imaging of SE-pHluorin-expressing neurons in anesthetized mice. A) Calibration curve constructed from ratiometric imaging of SE-pHluorin expressing hippocampal cultures in the presence of ionophores at a fixed Cl^-^ concentration of 5 mM. Mean and S.E.M. values are plotted. B) Representative maximal intensity projections of a vertically resliced data stack showing the SE-pHluorin signal and ratios at the somas of L2/3 pyramidal neurons from M1 cortex in the anesthetized state. Gray values were adjusted as follows: 530 nm, 710-15785; 480 nm, 187-970. C) Histogram showing the frequency distribution of the calculated values of pH_i_ obtained from the best fit calibration curve (A) and the average 530 nm/480 nm SE-pHluorin emission ratios from ROIs spanning the somas of L2/3 pyramidal neurons from M1 cortex in the anesthetized state.D) Average 530 nm/480 nm EGFP emission ratio from ROIs spanning the somas of EGFP expressing neurons as a function of depth of the soma centroid. Each color represents data from one out of three mice.

We sparsely expressed SE-pHluorin in superficial layers of motor cortex. Image stacks obtained from anesthetized mice revealed a similar compromise between the high intensity 530 nm emission and low intensity 480 nm emission as reported above for SCLM. In this case, the 480 nm emission at all processes including apical dendrites was too low to produce reliable ratios (double 480 nm emission than background fluorescence without saturating 530 nm emission). Consequently, we considered pyramidal cells having an oval/piriform soma with an extension representing the proximal portion of an apical dendrite. Thus, this *in vivo* imaging approach deep in intact brain tissue is limited to the somata of neurons and can not be used to assess pH_i_ in small compartments (Fig. 3B).

Using the best fit parameters we calculated the intracellular pH corresponding to the average ratios recorded *in vivo* at the somas of 27 L2/3 pyramidal neurons from 3 mice. We found pH_i_ values ranging from 6.5 to 7.5 with a mean of 7.1 ± 0.1 (mean ± 95% C.I., Fig. 3C).

To rule out ratio artefacts due to imaging depth and differential scattering of the emission wavelengths, we repeated our experiments employing EGFP, which has reduced pH sensitivity (pKa = 6, (Shaner et al., 2005)). We co-injected three mice with AAVs encoding Cre-dependent EGFP together with highly diluted Cre AAVs for bright and sparse labelling and repeated our measurements at L2/3 neurons using the same settings as with SE-pHluorin. We recorded somatic average emission ratios as a function of imaging depth. 3D stacks obtained from *in vivo* imaging sessions from 3 head fixed anesthetized WT mice revealed that the 530 nm/480 nm emission ratio of EGFP is not significantly correlated to the depth of the recorded soma in the tissue (Fig. 3D). Hence, depth-dependent scattering in the emission wavelengths of SE-pHluorin is unlikely to affect our *in vivo* ratiometric pH estimations.

Interestingly, the ratios recorded with SE-pHluorin, displayed a pronounced dispersion that was significantly greater than those recorded with EGFP (for average somatic ratios, standard deviation: EGFP = 1.8, n = 29 cells; SE-pHluorin = 2.5, n = 27 cells; p = 0.0010, Levene’s test). This suggests that the observed dispersion in SE-pHluorin ratios is indicating an actual variation in intracellular pH_i_ between cells *in vivo*.

### Calibration of SCLM and *in vivo* [Cl^-^]_i_ estimation

Ideally, pH_i_ and [Cl^-^]_;_ should be simultaneously deteR_min_ed in the same cellular compartment. However, SE-pHluorin imaging of neuronal pH_i_ is not compatible with simultaneous SCLM recordings. This means that it is not possible to calculate [Cl^-^]_;_ in a cell by cell basis from each cell’s SCLM ratio at the corresponding pH_i_. Hence, to estimate the average neuronal steady-state [Cl^-^]_i_ we constructed a calibration curve by clamping the [Cl^-^]_i_ and pH_i_ of SCLM-expressing hippocampal neurons to the extracellular saline through treatment with ionophores at a fixed pH corresponding to the average pH_i_ estimated *in vivo* (pH = 7.1, Fig. 4A). To match physiological conditions we constructed this curve at near physiological temperature (35 ± 2°C). In these conditions our best fit value of K_d_ for Cl^-^ was 13.6 ± 1.5 mM (r^2^ = 0.6972, Fig. 4 B). Using the best fit calibration curve, we estimated that the [Cl^-^]_i_ corresponding to the average SCLM ratio recorded *in vivo* at the somas of L2/3 pyramidal neurons of the motor cortex was 5 ± 2 mM in the anesthetized state (mean ± 95% CI).

**Figure 4:**
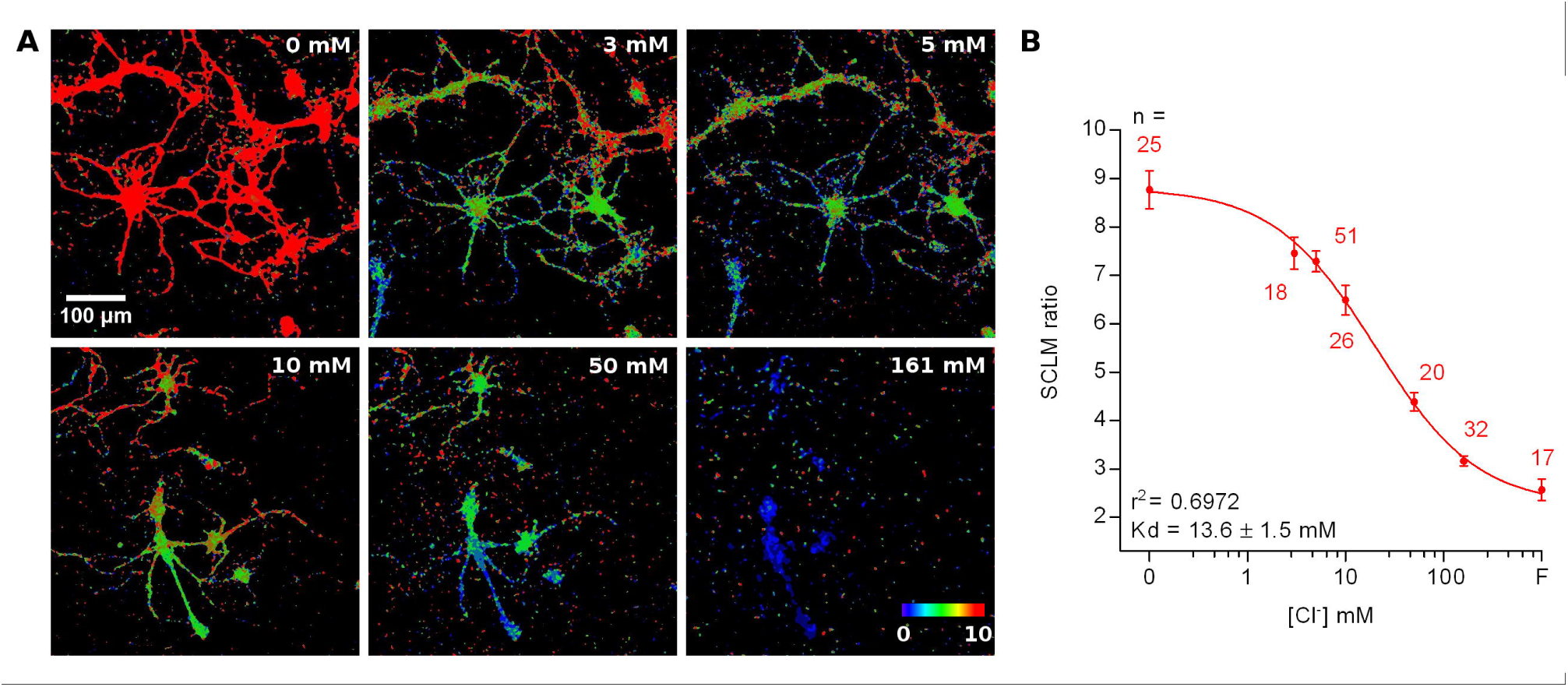
Calibration of SCLM. A) Representative images showing the *in vitro* SCLM ratio of hippocampal cultures in the presence of ionophores at different [Cl^-^] and a fixed pH of 7.1. To avoid deterioration of the cultures due to ionophore treatment, a maximum of 4 consecutive solutions were applied per coverslip. B) Calibration curve constructed from ratiometric imaging of SCLM expressing hippocampal cultures in the presence of ionophores at a fixed pH of 7.1. Mean and S.E.M. values are plotted. F = 153 mM fluoride solution.

### Other factors affecting ratiometric emission measurements and error estimates associated with pH changes

As mentioned above, the observed variability in our ratiometric measurements could be explained by irregularities in the optical properties of the tissue (e.g. presence of blood vessels) at different parts of the volume imaged. To explore this possibility we compared the Cerulean-Venus emission ratios obtained *in vivo* from L2/3 cortical neurons to the ones observed *in vitro* in hippocampal neuronal cultures expressing Cerulean-Venus. Thus, neuronal cultures should be significantly less influenced by uneven optical properties as the cells would be more exposed than they would when located deep in intact brain tissue. We observed that the dispersion of Cerulean-Venus expressing cells *in vitro* and *in vivo* did not differ significantly (for average somatic ratios, standard deviation: *in vivo* = 2.4, n = 52 cells; *in vitro* = 3.5, n = 18 cells; p = 0.1406, Levene’s test), indicating that the variability in our ratiometric measurements was not caused by tissue inhomogeneity (Fig. 5A), leaving pH_i_ differences between cells as a possible factor underlying this heterogeneity. To explore this further, we treated hippocampal cultures expressing Cerulean-Venus or SCLM with ionophores to manipulate their pH_i_ at a fixed [Cl^-^] of 5 mM and compared their ratios to our *in vivo* datasets (Fig. 5B). To compare the variability in the ratios from these *in vitro* recordings at different pH_i_ to our *in vivo* datasets we evaluated the ratio difference from the population mean ratio (signed ratio deviation, dRatio) at pH_i_ 7 (*in vitro* dataset) or 7.1 (*in vivo* dataset). We compared the span of the best fit curve and the 95% confidence band from the Cerulean-Venus or SCLM dRatio as a function of pH_i_ plots, within the range of pH_i_ estimated *in vivo* (6.5-7.5), to the ratio deviation obtained from *in vivo* measurements. In doing so, we observed that the pH sensitivity of Cerulean-Venus or SCLM could account for practically all of the variability observed *in vivo* (Fig. 5C, D). This result further supports that the observed variability in somatic SCLM ratios does not reflect a range of diverse neuronal [Cl^-^]_i_, but rather represents the range of pH_i_ we observed *in vivo*.

**Figure 5:**
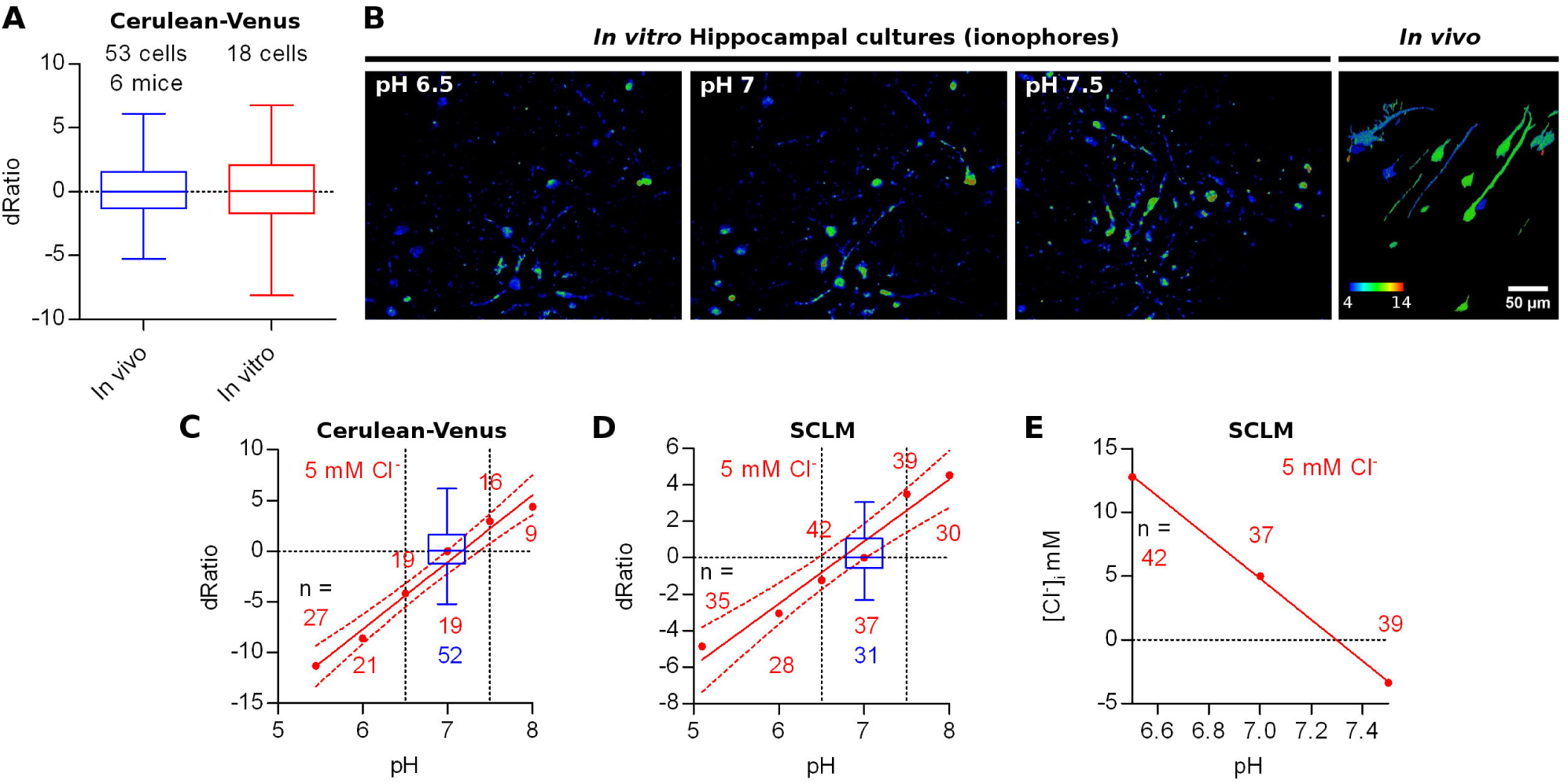
Other factors affecting ratiometric emission measurements. A) Ratio difference from the population mean ratio at pH 7-7.1 (signed deviation, dRatio) evaluated for Cerulean-Venus expressing L2/3 pyramidal neurons from M1 cortex (*in vivo*) or hippocampal cultures *(in vitro,* ACSF). B) Representative images showing the *in vitro* Cerulean-Venus ratio of hippocampal cultures in the presence of ionophores at different pH and a fixed [Cl^-^] of 5 mM (first 3 panels) and the *in vivo* Cerulean-Venus ratio variability (fourth panel). C) Red: Cerulean-Venus dRatio evaluated from hippocampal cultures in the presence of ionophores and different pH values. Mean dRatio and best fit linear regression plus 95% confidence band are plotted. Blue: dRatio distribution of Cerulean-Venus expressing L2/3 pyramidal neurons from M1 cortex *in vivo*. Segmented vertical lines: range of somatic pH measured *in vivo*. D) As in C but considering SCLM.E) Calculated [Cl^-^]_i_ values from the SCLM signals obtained in the experiment shown in D) at the *in vivo* pH_i_ range (6.5-7.5) using the calibration curve from Fig. 4 (constructed at pH 7.1).

Based on the quite pronounced variability of pH_i_ readouts, SCLM-expressing neurons may have a pH_i_ deviating from the average pH_i_ of 7.1, thereby introducing an error in the calculation of [Cl^-^]_i_ using the calibration curve constructed at the average *in vivo* pH_i_. We estimated the extent of error introduced by this approach by using the SCLM calibration curve produced at pH 7.1 to calculate the [Cl^-^]_i_ corresponding to the ratios measured *in vitro* for Figure 5D within the *in vivo* estimated pH_i_ range (Fig 3C). Within this pH range (6.5-7.5), if we use the SCLM calibration curve performed at pH 7.1 to calculate the [Cl^-^]_i_ in a cell by cell basis, an error of up to ± 8 mM could be introduced to the [Cl^-^]_;_ (Fig. 5E). Hence, this calibration approach using SCLM cannot accurately deteRmine the [Cl^-^]_i_ in a cell by cell basis but it can estimate the [Cl^-^]_i_ of a hypothetical average cell with average pH_i_. While the measurement error introduced with this approach will not suffice to resolve small changes in [Cl^-^]_i_, it might still be useful for experiments aimed at detecting larger changes in population [Cl^-^]_i_.

### *In vivo* imaging of SCLM and SE-pHluorin-expressing layer 2/3 cortical pyramidal neurons in awake mice

The anesthetized condition generally involves increased GABAergic inhibition (MacIver, 2014). This could influence [Cl^-^]_i_, producing estimates that are not representative of the physiological awake state. We imaged SCLM expressing neurons after initial isofluorane anesthesia in the awake state (Fig. 6A). We performed again a cell by cell pairwise comparison between the average somatic SCLM ratio and the average apical dendritic ratio of 24 L2/3 pyramidal cells from 5 mice and observed no significant difference (p = 0.5330, Wilcoxon signed rank test; Fig. 6B), suggesting that our previous observation about [Cl^-^]_i_ being homogeneous throughout these two subcellular compartments is not related to the anesthetized condition. Imaging in the awake state did not reveal significant differences in the distribution of somatic SCLM ratios (p = 0.9120, Kolmogorov-Smirnov; Fig. 6C), indicating that the steady-state neuronal [Cl^-^]_;_ is not affected by isoflurane anesthesia and that SCLM ratios measured under anesthesia are representative of the awake state. A cell by cell pairwise comparison between the average somatic SCLM ratio recorded from 24 identified L2/3 pyramidal cells from 5 mice in the anesthetized and awake state showed no significant difference (p = 0.6949, paired t-test; Fig. 6D), further supporting this notion.

**Figure 6:**
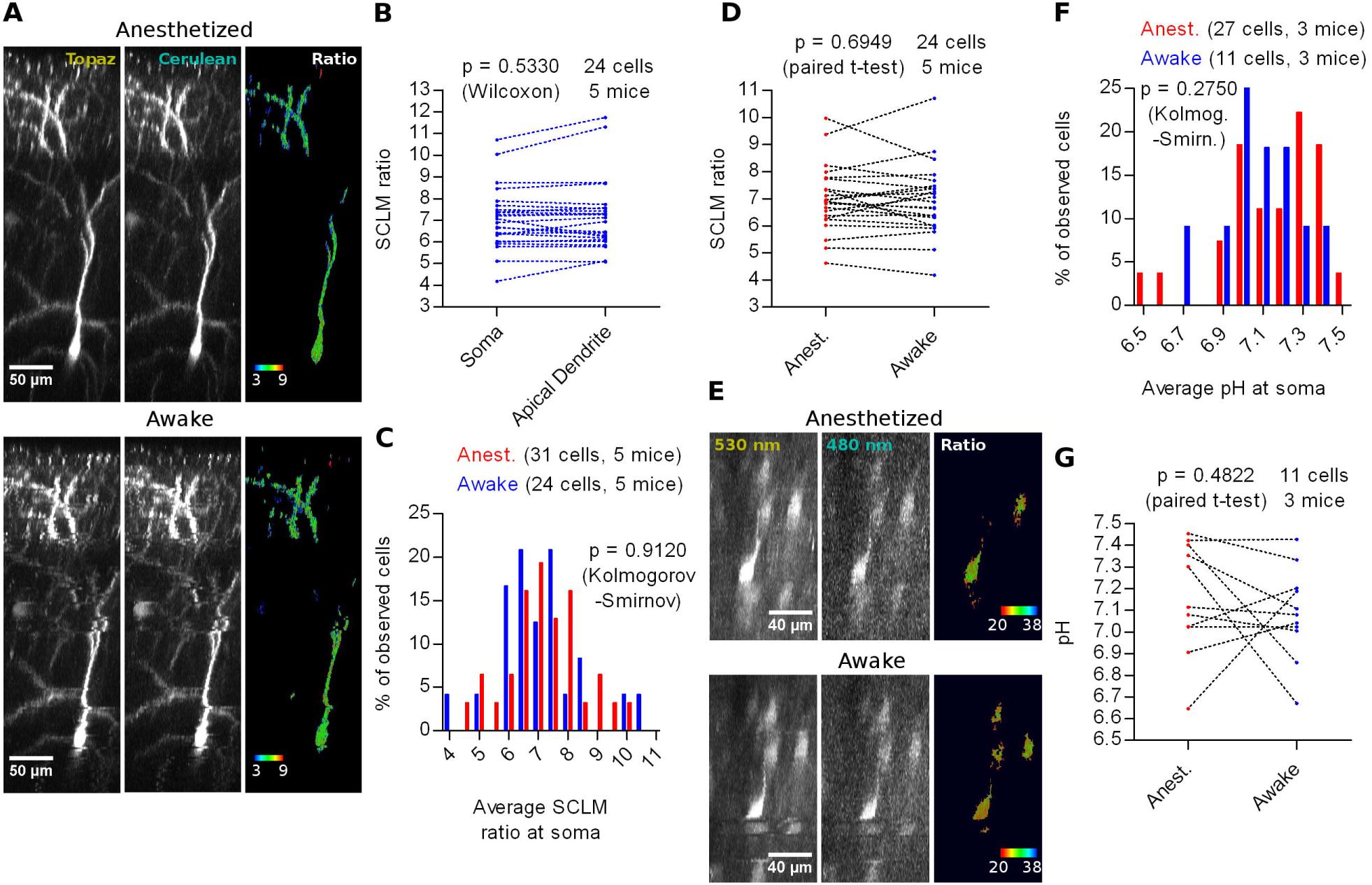
*In vivo* imaging of SCLM or SE-pHluorin-expressing neurons in awake mice. A) Representative maximal intensity projections of a vertically resliced data stack showing the *in vivo* SCLM Topaz and Cerulean signals and their corresponding ratio of a L2/3 pyramidal neuron from M1 cortex in the anesthetized and awake state. For display, gray values were clipped to include >90% of the total fluorescence signal: Topaz, 321-6240; Cerulean, 114-745. B) Pairwise comparison of the average SCLM ratio from ROIs spanning the basal apical dendrite and soma of L2/3 pyramidal neurons from M1 cortex in the awake state. C) Histogram showing the frequency distribution of the recorded average SCLM ratio from ROIs spanning the somas of L2/3 pyramidal neurons from M1 cortex in the anesthetized and awake states. D) Pairwise comparison of the average SCLM ratio from ROIs spanning the soma of L2/3 pyramidal neurons from M1 cortex in the anesthetized and awake state. E) Representative maximal intensity projections of a vertically resliced data stack showing the SE-pHluorin signal and ratios at the somas of L2/3 pyramidal neurons from M1 cortex in the anesthetized and awake state. Gray values were adjusted as follows: 530 nm, 899-11836; 480 nm, 168-695. F) Histogram showing the frequency distribution of the calculated values of pH_i_ obtained from the calibration curve and the average 530 nm/480 nm SE-pHluorin emission ratios from ROIs spanning the somas of L2/3 pyramidal neurons from M1 cortex in the anesthetized and awake state. G) Pairwise comparison of the somatic pH of L2/3 pyramidal neurons from M1 cortex in the anesthetized and awake state.

To estimate [Cl^-^]_i_ in the awake state using a calibration curve, we first studied pH_i_. Hence, we imaged SE-pHluorin in the awake state (Fig. 6E) and found that the pH_i_ distribution is not significantly different to the one obtained under isoflurane anesthesia (p = 0.2750, Kolmogorov-Smirnov; Fig. 6F). This result suggests that imaging SE-pHluorin under isoflurane anesthesia also does not produce artefactual measures of steady-state pH_i_. A cell by cell pairwise comparison between the average somatic pH recorded from 11 identified L2/3 pyramidal cells from 3 mice in the anesthetized and awake state showed no significant difference (p = 0.4822, paired t-test; Fig. 6G).

More importantly, these results suggest that the previous observation about the steady-state SCLM ratio being unaffected by anesthesia is truly reflecting a lack of effect on [Cl^-^]_i_. Using our best fit calibration curve (Fig. 5), we estimated the [Cl^-^]_i_ using the average somatic SCLM ratio recorded *in vivo* in the awake state to be 6 ± 2 mM (mean ± 95% C.I.).

To conclude, we did not find differences in [Cl^-^]_;_ or pH_i_ in cortical neurons of isofluorane anesthetized versus awake mice, suggesting that steady-state [Cl^-^]_i_ or pH_i_ are not affected by isoflurane anesthesia.

### *In vivo* effect of KCC2 deletion on the steady-state [Cl^-^]_i_ of adult layer 2/3 cortex neurons

As a proof of principle of our *in vivo* imaging approach and to explore the factors affecting the steady-state neuronal intracellular Cl^-^ **in vivo*,* we co-injected AAVs encoding Cre-dependent SCLM together with highly diluted AAVs encoding Cre recombinase into L2/3 of M1 cortex in adult KCC2^lox/lox^ conditional KO mice (Seja et al., 2012; Godde et al., 2016). Thus, expression of Cre recombinase will yield SCLM expression and abolish expression of KCC2. These mice were imaged at least 4 weeks after virus injection, which widely exceeds the reported KCC2 protein turnover time (Rivera et al., 2004). As a control for [Cl^-^]_i_ estimations, we injected AAVs encoding Cre-independent SCLM to assess [Cl^-^]_i_ in neurons of KCC2^lox/lox^ mice without inducing the KO (from now on Ctrl cells). Additionally, to validate these measurements we determined the pH_i_ in Ctrl cells and KCC2 KO cells of KCC2^lox/lox^ mice using SE-pHluorin and following the aforementioned approach (+/-Cre). The expression of Cre-independent SCLM and SE-pHluorin under the Synapsin promoter *in vivo* did not permit to distinguish pyramidal cell morphology as clearly as our Cre-dependent dilution approach, as the concomitant labelling of the tissue is much denser. Therefore, in these cases cells with clear pyramidal neuron morphology were carefully selected for further analysis, thus discarding the great majority of labelled cells.

*In vivo* imaging of SE-pHluorin labelled KCC2 KO and Ctrl cells from 6 KCC2^lox/lox^ mice showed that the KO did not affect the pH_i_ distribution (Anesthetized: p = 0.0990; Awake: p = 0.2980; Kolmogorov-Smirnov; Fig. 7A). Again, we confirmed in the kCC2^lox/lox^ genetic background, that isoflurane anesthesia does not affect the steady-state neuronal pH_i_ distribution (+ Cre: p = 0.8180; - Cre: p = 0.2710; Kolmogorov-Smirnov; Fig. 7A).

**Figure 7:**
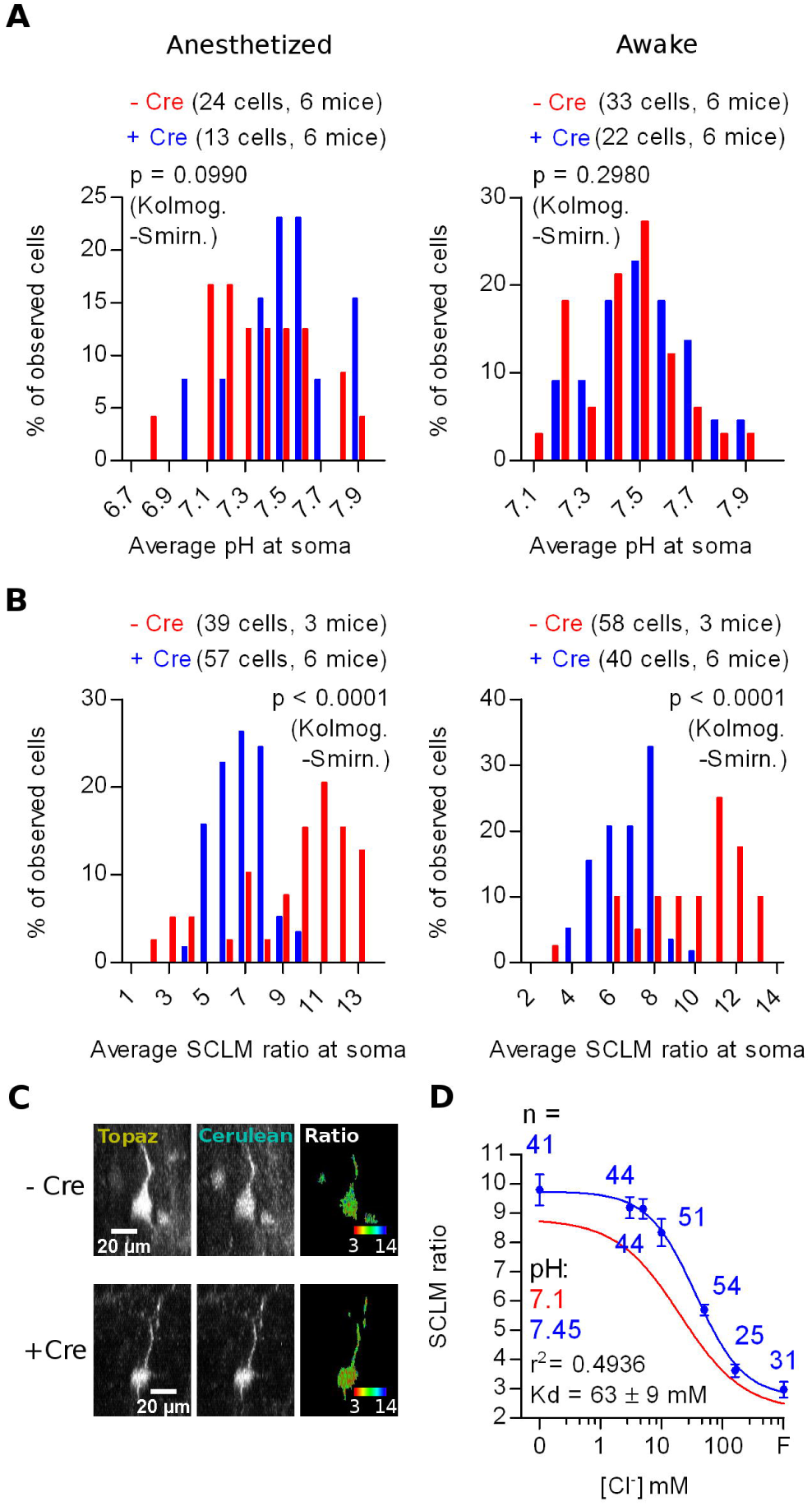
*In vivo* effect of KCC2 deletion on the steady-state [Cl^-^], of adult layer 2/3 cortex neurons. A) Histogram showing the frequency distribution of the calculated values of pH_i_ obtained from *in vivo* recordings in anesthetized and awake KCC2^lox/lox^ mice through Cre-dependent (KO for KCC2) and Cre-independent SE-pHluorin expression. B) Histogram showing the frequency distribution of the average SCLM 540 nm/480 nm emission ratio from ROIs spanning the somas of L2/3 pyramidal neurons from M1 cortex in KCC2^lox/lox^ mice through Cre-dependent (KO for KCC2) and Cre-independent SCLM expression recorded in the anesthetized and awake state (left and right panels respectively). C) Representative maximal intensity projections showing the Cre-dependent and Cre-independent SCLM signal and ratios at the soma of a L2/3 pyramidal neuron from M1 cortex of KCC2^lox/lox^ mice in the anesthetized state. Gray values were adjusted for display as follows: Topaz - Cre, 1061-15758; Cerulean - Cre, 176-1700; Topaz + Cre, 744-13895; Cerulean + Cre, 143-1717. For easier visualization, the ratiometric look-up table is inverted with respect to the other ratiometric SCLM panels to highlight the shift towards higher [Cl^-^]_i_ as a shift towards red instead of a shift towards blue in a dark background. D) Calibration curve constructed from ratiometric imaging of SCLM expressing hippocampal cultures in the presence of ionophores at a pH of 7.45 (blue). Mean and S.E.M. values are plotted. The best fit calibration curve obtained at pH 7.1 is plotted in red for comparison, informed parameters correspond to the blue calibration curve.

*In vivo* imaging of SCLM-labelled KCC2 KO cells from 6 mice showed significantly different average somatic SCLM ratio distributions than Ctrl (- Cre) cells from 3 mice (Anesthetized: p < 0.0001; Awake: p < 0.0001; Kolmogorov-Smirnov; Fig. 7B, C). Once again, we observed no significant effect of isoflurane anesthesia in the SCLM ratio distributions from KCC2^lox/lox^ mice, further supporting that isoflurane does not affect the distribution of steady-state neuronal [Cl^-^]_;_ (+ Cre: p = 0.6650; - Cre: p = 0.8600; Kolmogorov-Smirnov; Fig. 7B). The observed lack of change in the pH_i_ distribution (Fig. 7A) confirms that the observed changes in the SCLM ratio distribution induced by deleting KCC2 (Fig. 7B) are caused by changes in the steady-state neuronal [Cl^-^]_;_, ruling out the influence of pH on SCLM ratios. Altogether, these results point out that the KO of KCC2 produces a shift towards lower SCLM ratios, which represent higher neuronal [Cl^-^]_i_ and that this effect is discernible *in vivo* using SCLM.

To calculate an estimate of the change in the average neuronal steady-state [Cl^-^]_i_ we constructed an *in vitro* calibration curve as previously described. We chose to produce this calibration curve at pH 7.45 because this value was not significantly different to the mean pH_i_ recorded in KCC2 KO cells from KCC2^lox/lox^ mice in both anesthetized or awake conditions (Anesthetized: p = 0.2163; Awake: p = 0.1941; Wilcoxon signed rank test). Thus, we used this single calibration curve produced at pH 7.45 to calculate estimates of the steady-state [Cl^-^]_i_ from the mean SCLM ratios recorded *in vivo* at the somas of KCC2 KO cells. This calibration curve showed a higher maximum ratio and K_d_ than the one produced at pH 7.1 (Fig. 7D). The resulting steady-state [Cl^-^]_i_ estimation produced by KCC2 KO was 24.2 ± 8 mM in the anesthetized state and 25.7 ± 8 mM in the awake state (mean ± 95% C.I.). In summary, KCC2 deletion produced a higher neuronal [Cl^-^]_;_, demonstrating a requirement of KCC2 for low [Cl^-^]_i_ in the adult mouse brain. Furthermore, These results indicate that effects in the magnitude of the changes reported here can be detected by *in vivo* SCLM imaging.

## Discussion

Our study reports non-invasive *in vivo* steady-state [Cl^-^]_i_ and pH_i_ estimations in L2/3 cortical neurons of anesthetized and awake mice. Independently of anesthesia, we estimated a native steady-state neuronal [Cl^-^]_i_ of 6 ± 2 mM which is consistent with numerous *in vitro* reports (Berglund et al., 2008; Bregestovski et al., 2009; Raimondo et al., 2013). According to the Nernst equation, a 6 mM steady-state neuronal [Cl^-^]_i_ paired with the expected extracellular [Cl^-^] (145 mM) (Raimondo et al., 2015) would produce an E_C1_ of ∼(-85 mV) at 37°C, which is compatible with the expected values of healthy adult neurons (Doyon et al., 2016). If the Goldman equation is used to calculate the respective GABA_A_R E_rev_ using the reported [Cl^-^]_i_ and the expected [HCO_3_^-^]; (15 mM) at the reported average pH_i_, paired with the expected extracellular [Cl^-^] (145 mM) and [HCO_3_^-^] (24 mM) concentrations (Raimondo et al., 2015) and a 4:1 permeability ratio for these two anions (Bormann et al., 1987), an E_rev_ of ∼(-73 mV) is obtained, consistent with previously reported values (Kaila et al., 1993; Lee et al., 2015). This GABA_A_ E_rev_ could support hyperpolarizing inhibition, although the driving force at typical resting membrane potentials would be rather small. Therefore, Cl^-^ conductances will tend to produce shunting inhibition **in vivo*.* Hence our results demonstrate that somatic steady-state [Cl^-^]_i_ deteR_min_ed *in vitro* can, in principle, be extrapolated to the native state found in the awake brain.

In this study we did not find evidence of differences in neuronal [Cl^-^]_i_ along the soma to proximal regions of the apical dendrite of cortical L2/3 pyramidal neurons. This is in contrast to *in vitro* reports studying other cell types such as cultured hippocampal or spinal cord neurons and retinal bipolar cells where clear somato-dendritic gradients were detected (Kuner and Augustine, 2000; Duebel et al., 2006; Waseem et al., 2010). Thus, our study suggests that within these subcellular compartments GABAergic inputs would be homogeneously inhibitory. The work of Waseem et al. (2010) in cultured neurons reports a stark gradient in [Cl^-^]_;_ between the soma and proximal dendritic compartment within a ∼20 μm distance. Nevertheless, this observation can be influenced by the maturation stage of the cultured cells, limited neuron-glia interactions and other uncontrolled factors inherent to this *in vitro* preparation such as abnormal neuronal activity, which can affect cation-Cl^-^ co-transporter trafficking and activation (Kaila et al., 2014). In this respect our study undertakes the effort to keep the intact native conditions of brain tissue as close as possible to the physiological state to avoid these issues. Under these conditions, we did not find evidence of a [Cl^-^]_i_ gradient at the proximal portion (up to ∼100 μm away from the soma) of the apical dendrite of L2/3 pyramidal neurons.

We found a steady-state pH_i_ of 7.1 ± 0.1 mM at the neuronal soma which is consistent with *in vitro* reports (Caspers and Speckmann, 1972; Raimondo et al., 2012). Our *in vivo* estimations of neuronal pH_i_ suggest that individual L2/3 pyramidal neurons differ in pH_i_ across a large range, also consistent with *in vitro* reports (Schwiening and Boron, 1994; Ruffin et al., 2014) and a recent *in vivo* report (Sulis Sato et al., 2017). Neuronal activity can dynamically affect pH_i_ (Chesler and Kaila, 1992). Thus it is possible that cells with different activity levels would underlie the observed range in pH_i_. We could not systematically explore the occurrence of gradients in pH_i_ along the somato-dendritic axis, which is necessary to validate our result suggesting homogeneous [Cl^-^]_i_ along this compartment. Nevertheless, the occurrence of simultaneous [Cl^-^]_i_ and pH_i_ gradients producing perfectly opposing effects on SCLM ratios is highly unlikely. This is supported by our observation that somato-dendritic pH gradients could not be detected with the Cl^-^-insensitive but slightly pH-sensitive Cerulean-Venus protein that otherwise reported cell-to-cell differences in pH. In conclusion we found that under native conditions of the awake adult brain, pH_i_ differs between cells by a large margin.

We provide *in vivo* evidence of the essential role of KCC2 in producing low neuronal [Cl^-^]_i_ in adult mouse neurons (Kaila et al., 2014; Doyon et al., 2016). In doing so, we show that despite the limitations of the SCLM/SE-pHluorin based *in vivo* imaging approach, it could resolve changes in neuronal [Cl^-^]_;_ that would be compatible to the ones expected during the developmental shift in [Cl]; (Ben-Ari, 2002; Kaila et al., 2014). Thus, a plausible future perspective of this work would be to explore the developmental [Cl^-^]_;_ shift in different genetically defined neuronal populations.The KO of KCC2 is expected to impact neuronal activity as it is evidenced in reports using pharmacological inhibitors of KCC2 in primary neuronal cultures, acute brain slices and *in vivo* (Sivakumaran et al., 2015). In consequence, this change in neuronal activity could affect pH_i_ (Chesler and Kaila, 1992). Nevertheless, the sparse AAV-driven Cre recombinase expression in KCC2^lox/lox^ mice affect only a small subset of neurons in the cortex in contrast to the global network effect of pharmacological KCC2 inhibitors which leads to epileptiform activity. Hence, even though the Cl^-^ driven synaptic inputs to KCC2 KO neurons would likely cause excitation, the overall inputs to the neuron would not be as drastically enhanced as in an epileptiform event, consistent with the observed lack of effect of the KCC2 KO on pH_i_. Nonetheless, further controls of neuronal activity in KCC2 KO and Ctrl cells should be produced to fully address this point in future research.

Finally, our study provides a foundation for future *in vivo* research employing Cl^-^ and pH imaging in healthy and diseased tissue. While this paper was under review, Sulis Sato et al. (2017) published an *in vivo* Cl^-^ and pH imaging study using ClopHensor. In the following paragraphs we briefly compare the results produced by both sensors and discuss their strengths and caveats. *in vivo* 2P imaging of SCLM or SE-pHluorin showed a modest subcellular resolution. The main factor limiting this is the restricted 2P illumination settings needed to excite SCLM without saturating the Topaz emission channel, still obtaining reliable signal to noise levels in the Cerulean channel. Shorter 2P excitation wavelengths (800 nm) did not significantly improve this (data not shown). Alternatively, changing the relative sensitivity of the PMTs or replacing Cerulean for the higher quantum yield variant Teal, dramatically reduced the dynamic range of SCLM, thus practically abolishing the ability to resolve different [Cl^-^]_i_ (data not shown). Hence, the settings reported here represent the best compromise we found, still allowing us to explore the subcellular steady-state [Cl^-^]_;_ between the soma and apical dendrite of pyramidal neurons. Sulis Sato et al. (2017) did not explore subcellular differences in [Cl^-^]_;_ and pH_i_ due to better signal-to-noise levels at the soma compared to the dendrites, implying a similar limitation of ClopHensor.

The approach we describe to produce ratiometric Cl^-^ and pH estimations is not affected by isoflurane anesthesia. The anesthetized condition generally involves increased GABAergic inhibition (MacIver, 2014) and can have profound effects on breathing which could potentially alter cellular pH homeostasis (Massey et al., 2015). In the case of isoflurane, it has been reported to significantly enhance GABAergic inhibition in rat hippocampal slices and to mask the chemosensitivity of pH/CO2 sensitive serotonergic neurons of the medulla (MacIver, 2014; Massey et al., 2015). Despite these known effects of isoflurane, *in vivo* optical steady-state estimations of [Cl^-^]_i_ and pH_i_ in L2/3 pyramidal neurons using SCLM and SE-pHluorin are not significantly affected by this anesthetic.The study of Sulis Sato et al. (2017) found similar [Cl^-^]_;_ and pH_i_ under urethane anesthesia.

Our approach was also not affected by light scattering caused by tissue depth or uneven optical properties. The use of ratiometric sensors such as SCLM deep in scattering tissue is expected to be hampered by differential scattering of two spectrally different fluorescence emissions. This study was started with such expectation in mind and fluorescence life time imaging (FLIM) of the SCLM donor (Cerulean) was employed to obtain a depth-independent readout. However, the changes in life time as a function of [Cl^-^] in *in vitro* calibration curves were too small in order to resolve changes of [Cl^-^] in the physiologically relevant range (data not shown). The emission ratio of an environmentally less sensitive construct, Cerulean-Venus, did not significantly change up to a depth of approximately 350-400 μm. Therefore, motor cortex, and possibly brain tissue in general, appears to scatter emission light in the range from 480 to 540 nm in a manner so similar that changes in the ratio as a function of depth were not evident, suggesting that this spectral range can be used for ratiometric recordings without applying complicated depth-dependent corrections. This is reported to be fundamentally different when ratiometric indicators using the green-red range of the emission spectrum are employed, such as ClopHensor, as brain tissue shows a strong depth-dependent differential scattering for this range of the spectrum (Sulis Sato et al., 2017). On the other hand excitation light of different wavelengths can be differentially scattered as a function of imaging depth producing “distortions” on the absorption spectra of fluorescent proteins (Sulis Sato et al., 2017). Nevertheless, our ratiometric imaging approach required only one excitation wavelength, making it relatively less prone to be affected by this issue in comparison to ClopHensor (Sulis Sato et al., 2017). Again, this is supported by the observed lack of correlation between the imaging depth and the recorded ratios for Cerulean-Venus or EGFP expressing neurons *in vivo*.

Additionally the presence of blood vessels and fiber bundles can make the optical properties of brain tissue inhomogeneous, which can also cause scattering artefacts. By comparing the Cerulean-Venus ratios recorded *in vivo* to the ones recorded in hippocampal cultures we demonstrated that our ratiometric approach is not significantly affected by this factor. Altogether, we provide evidence supporting that ratiometric imaging in the emission range from 480 to 540 nm is robust against these inherent scattering-related issues of ratiometric imaging in intact brain tissue, an insight that will be relevant for *in vivo* applications of a multitude of genetically encoded indicators employing the CFP-YFP FRET pair.

The use of SCLM for intracellular Cl^-^ estimations needs validation due to its pH sensitivity. Yet, simultaneous recordings of pH_i_ and [Cl^-^]_;_ using this approach are technically not yet possible. The observed diverse range of neuronal pH_i_ makes SCLM readouts quite variable between individual cells. This is a major limitation in SCLM imaging that demands a compatible method to simultaneously measure pH_i_ and achieve calculations of [Cl^-^]_i_ for individual cells. Additionally, as neuronal activity also affects pH_i_ (Chesler and Kaila, 1992), this also stresses the need of simultaneous pH_i_ readouts to assess dynamic neuronal Cl^-^ changes using SCLM.

In conclusion, this report contributes to the study of neuronal [Cl^-^]_i_ and pH_i_ by providing estimations as close to the native state as practically possible. These results and approach represent a step forward towards the understanding of essential neuronal processes involving [Cl^-^]_i_ in the intact organism and aid in the quest to relieve pathophysiological conditions such as epilepsy or schizophrenia (Kaila et al., 2014; Sullivan et al., 2015).

## Non-standard abbreviations

SCLM: Superclomeleon.
SE-pHluorin: Superecliptic pHluorin.

## Author Contributions

JB, JK and TK designed research, reviewed and edited the manuscript. MK designed and constructed expression plasmids. JB and JK performed imaging experiments. JB performed data analysis and prepared figures. JB and TK wrote the manuscript.

## Additional Information

The authors declare no competing financial or commercial interests.

## Acknowledgements

TK acknowledges funding by the CellNetworks Cluster of Excellence (EXC 81) and a large equipment grant by the DFG. We thank Prof. Dr. Thomas Jentsch, Prof. Dr. George Augustine, Prof. Dr. Gero Miesenböck, Prof. Dr. Dave Piston, Dr. Atsushi Miyawaki, Prof. Dr. Jochen Kuhse and Prof. Dr. Hilmar Bading for providing materials. We would also like to thank Dr. Christoph Körber and Dr. Patric Pelzer for helpful discussion and comments on the manuscript, Claudia Kocksch and Gabriela E. Krämer for technical assistance. The authors gratefully acknowledge the data storage service SDS@hd supported by the Ministry of Science, Research and the Arts Baden-Württemberg (MWK) and the German Research Foundation (DFG) through grant INST 35/1314-1 FUGG. We acknowledge the financial support of the Deutsche Forschungsgemeinschaft and Ruprecht-Karls-Universität Heidelberg within the funding programme Open Access Publishing.

